# Diverse domain architectures of CheA histidine kinase, a central component of bacterial and archaeal chemosensory systems

**DOI:** 10.1101/2023.09.19.558490

**Authors:** Marissa A. Berry, Ekaterina P. Andrianova, Igor B. Zhulin

## Abstract

Chemosensory systems in bacteria and archaea are complex, multi-protein pathways that enable rapid cellular responses to environmental changes. The CheA histidine kinase is a central component of chemosensory systems. In contrast to other histidine kinases, it lacks a sensor (input) domain and utilizes dedicated chemoreceptors for sensing. CheA is a multi-domain protein; in model organisms as diverse as *Escherichia coli* and *Bacillus subtilis*, it contains five single-copy domains. Deviations from this canonical domain architecture have been reported, however, a broad genome-wide analysis of CheA diversity is lacking. Here, we present results of a genomic survey of CheA domain composition carried out using an unbiased set of thousands of CheA sequences from bacteria and archaea. We found that four out of five canonical CheA domains comprise a minimal functional unit (core domains), as they are present in all surveyed CheA homologs. The most common deviations from a classical five-domain CheA architecture are the lack of a P2/CheY-binding domain, which is missing from more than a half of CheA homologs and the acquisition of a response regulator receiver (CheY-like) domain, which is present in ∼35% of CheA homologs. We also document other deviations from classical CheA architecture, including bipartite CheA proteins, domain duplications and fusions, and reveal that phylogenetically defined CheA classes have pre-dominant domain architectures. This study lays a foundation for a better classification of CheA homologs and identifies targets for experimental investigations.

## Introduction

Chemosensory systems (also often called chemotaxis systems) are multi-protein pathways that enable fast “decision making” in bacteria and archaea in response to rapid changes in their microenvironment, such as increasing or decreasing concentrations of nutrients and toxins (1, 2). A pathway regulating chemotaxis in *Escherichia coli* is the best studied chemosensory system (3, 4). The histidine kinase CheA (5) is the central component of this system. CheA activity is modulated by chemoreceptors, also called methyl-accepting chemotaxis proteins (MCPs), that sense external and intracellular signals (3). Chemoreceptors, CheA and the CheW adaptor protein form signaling complexes that assemble into chemosensory arrays (6, 7). An increase in repellent or decrease in attractant concentrations increases CheA autophosphorylation and phosphorylated CheA donates the phosphate group to the response regulator protein CheY (8). Phosphorylated CheY binds to flagellar motors and promote clockwise rotation of the flagella and cell tumbling. When CheA activity is reduced by attractants, a decrease in concentration of phosphorylated CheY promotes counterclockwise rotation of the flagella and smooth swimming. The CheR methyltransferase and the CheB methylesterase that covalently modify chemoreceptors comprise an adaptation pathway (9). The *E. coli* system also includes the CheZ phosphatase, which de-phosphorylates CheY resulting in signal termination (10).

Chemosensory systems homologous to the *E. coli* pathway were identified and experimentally studied in other bacterial and archaeal species, where they were shown to control not only flagellar motility, but also type IV pili (Tfp) based motility, biofilm formation, cell-cell interaction, biosynthesis, development, and other cellular functions (11–14). By regulating these vital processes, chemosensory systems ultimately have a strong impact on bacterial behavior, lifestyle, and interactions with hosts and between species (15, 16). Comparative genomic analysis suggested that approximately half of bacterial species contain chemosensory pathways (17). In terms of component design, chemosensory systems are the most complex mode of signal transduction in bacteria (18). Although the number of components vary organism to organism and system to system, four core proteins are present in every system: chemoreceptors, CheA, CheW, and CheY (17). CheA is an ideal marker for studying the diversity of chemosensory systems because (i) it is a large, multi-domain protein (5), (ii) it is present in every system (17), and (iii) there is always only one copy of CheA per system, which is not the case for other three core proteins (17).

The current knowledge on the structure and function of CheA primarily comes from work on *Escherichia coli* and *Thermotoga maritima* (19) that are model organisms for functional and structural studies, respectively. CheA proteins from both organisms consist of five domains, that were initially labeled P1 through P5 (20) and were later recognized as members of conserved domain families (21) (Fig. 1, Table S1). The leading protein domain database Pfam (22), which is now a part of the InterPro resource (23), contains profile models for the five canonical CheA domains, which show their relationship with other protein families and allow for their identification in genomic datasets (Table S1).

**Fig. 1.**
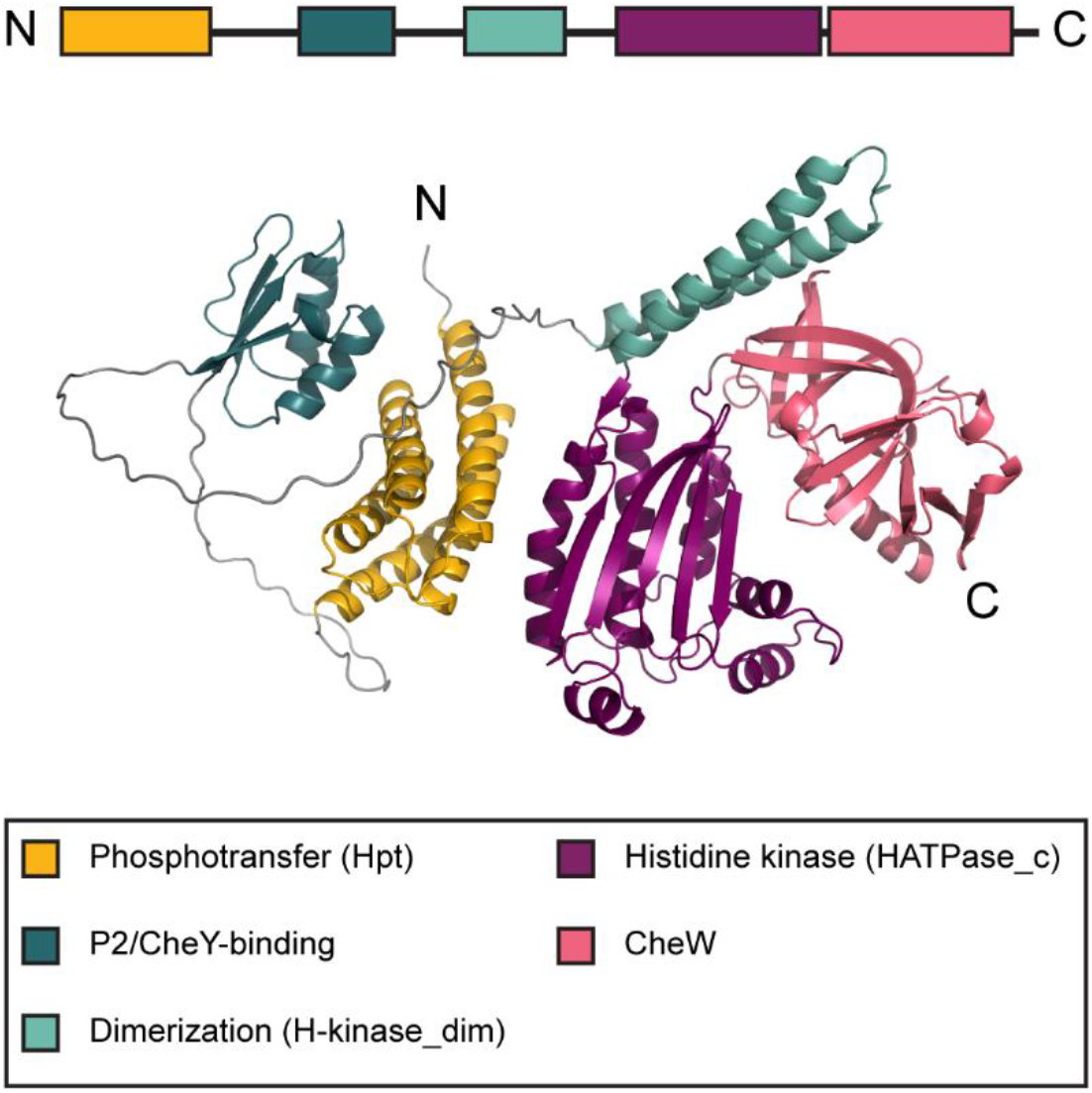
Classical CheA domain architecture. Domain composition of the CheA protein from *E. coli* (accession number WP_001350517.1). Structure of the full-length protein was predicted by AlphaFold (48).

N-terminal P1 is the **<u>h</u>**istidine-containing **<u>p</u>**hosphotransfer domain (Hpt); it is responsible for the movement of the phosphate group from its substrate histidine residue to CheY and CheB response regulators (24, 25). P2 is a docking domain, which binds CheY and CheB (20, 26). It is not required for phosphotransfer, but it greatly accelerates its rate (20). The P3 **<u>dim</u>**erization domain (H-kinase_dim) mediates CheA dimer formation (5). The P4 histidine kinase domain (HATPase_c: **h**istidine kinase-, DNA gyrase B-, and HSP90-like **ATPase**) binds ATP (27). The C-terminal P5 domain (CheW) is homologous to CheW protein (5, 21); it couples CheA to CheW and chemoreceptors enabling the formation of a chemosensory signaling complex (28). The leading protein domain database Pfam (22), which is now a part of the InterPro resource (23), contains profile models for the five canonical CheA domains, which show their relationship with other protein families and allow for their identification in genomic datasets (Table S1).

A phylogenomic study classified CheA proteins in more than a dozen of classes based on sequence similarity and genomic context (17). Most of these classes were predicted to govern **f**lagellar motility and were termed F1 through F17, whereas two classes were predicted to control **<u>T</u>**ype **<u>IV p</u>**ili based motility (termed Tfp class) and other (**<u>a</u>**lternative) **<u>c</u>**ellular **f**unctions (termed ACF class) (17). In that study, a comparative CheA analysis was perform using the unit of three domains – dimerization (H-kinase_dim), histidine kinase (HATPase_c) and CheW – because they were detected in all CheA sequences and were always in the same configuration, which was not the case for the other two domains, Hpt and P2/CheY-binding (17). Interestingly, structural studies also resulted in defining the last three domains as a core unit (5), whereas structures of the Hpt domain and P2/CheY-binding domain were solved separately because these domains are separated by long linker regions in both *E. coli* and *T. maritima* (26). This five domain arrangement is considered as the canonical domain architecture of CheA, because it was reported in key model organisms including not only *E. coli* and *T. maritima*, but also *Salmonella enterica* (29), *Bacillus subtilis* (30) and an archaeon *Halobacterium salinarum* (13). However, subsequent studies with other bacterial species revealed some variation in CheA domain composition. For example, CheA of *Helicobacter pylori* and *Campylobacter jejuni* do not contain P2/CheY-binding domain, but have a response regulator receiver domain in their C-terminus (31). *Azospirillum brasilense* and *Azorhizobium caulinodans* CheA proteins contain two CheW domains and a response regulator receiver domain (32, 33). CheA of Tfp class in *Pseudomonas aeruginosa* has a response regulator receiver domain in the C-terminus, lacks P2/CheY-binding domain and has multiple Hpt domains in the N-terminus (34). Although these findings revealed substantial and intriguing variations in the CheA domain composition and arrangement, the extent of this diversity is unknown. In this study, we aimed to fill this gap by projecting our current understanding of CheA structure and function onto the current genomic landscape.

## Results and Discussion

### Constructing a representative set of CheA protein sequences

The number of microbial genomes is rapidly increasing, and currently reaches almost 1.4 million of bacterial genomes in the NCBI database (35). However, the impressive amount of the sequenced genomes does not reflect the phylogenetic diversity, as 90% of these genomes belong to a very few (out of more than a hundred) phyla, thus, making this dataset heavily biased and leaving most bacterial phyla highly underrepresented. Therefore, to address this bias, for our CheA search we decided to use a much more balanced set of representative genomes from the Genomic Taxonomy database (GTDB) (36). The current version GTDB v.95 contains a representative set of 16859 RefSeq (37) genomes spanning 50 bacterial and 18 archaeal phyla. For rapid and convenient CheA identification and classification we restricted the dataset to the genomes that were available in the MIST 3.0 database (38), resulting in 14796 genomes. Among those genomes, 7900 did not have any identifiable CheA protein. Available archaeal genomes are not as numerous, thus we collected CheA sequences from all archaeal genomes available at GTDB (see Materials and Methods for details). Our final dataset contained 13673 CheA protein sequences from 6896 bacterial and 471 archaeal genomes, that represent 36 bacterial and 8 archaeal phyla and all CheA classes: F1 through F17, ACF and Tfp (Fig 2). The domain composition was then determined for all collected CheA sequences and analyzed with respect to domain presence, absence, and duplication. A complete list of analyzed CheA sequences along with genome, taxonomy, domain composition, and other relevant data can be found in a GitHub repository (https://github.com/mberry14/Diversity_of_CheA_domain_architectures).

**Fig. 2.**
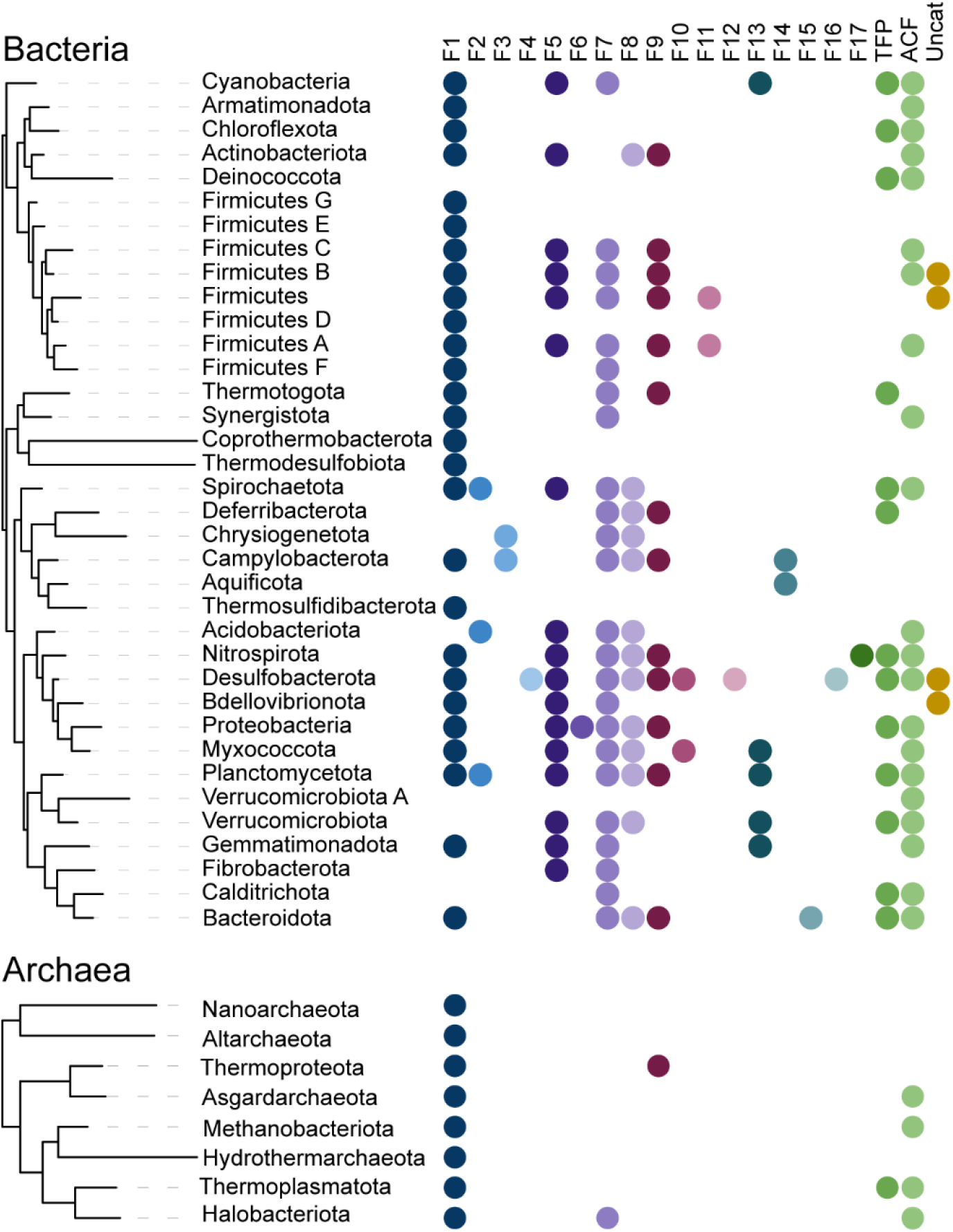
Phyletic distribution and phylogenomic classes of CheA proteins analyzed in this study. Phylogenetic tree based on 120 most common genes was adopted from AnnoTree (59). Phylogenomic classes are defined in (17).

### Core domains of CheA

Our analysis revealed substantial variations in CheA domain composition: most frequently found domain combinations are shown in Fig. 3A. Unexpectedly, we found that only 46% of the CheA protein sequences in our dataset have the classical five domain architecture (Fig. 3). The most common deviation from this paradigm was the lack of P2/CheY-binding domain - nearly 52% of all the CheA sequences are missing this domain (Fig. 3B). Consequently, we define the core CheA domains (present in all CheA sequences and therefore define its function).

**Fig. 3.**
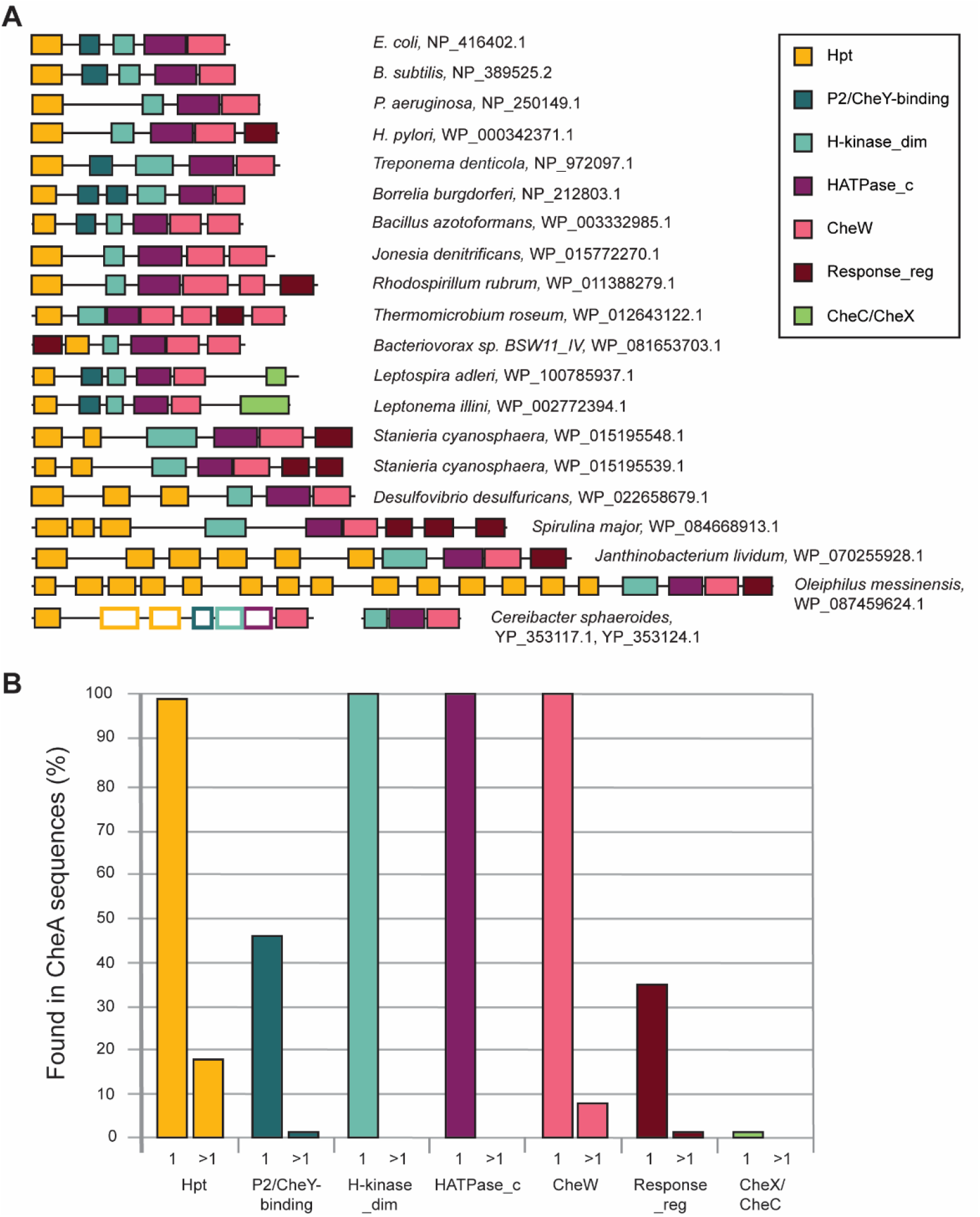
CheA domain composition. **A)** Representative domain architectures; species names and accession numbers for CheA proteins are shown. Domains detected by structural but not sequence similarity are shown as empty rectangles. **B)** Core and auxiliary domains and their duplications. Percentage of CheA sequences with a given domain is shown. Single domain occurrence is labeled as 1 on the x axis; >1 depicts sequences with more than one copy of a given domain.

#### The histidine kinase domain (HATPase_c)

As expected, this domain is present in every CheA (Fig. 3B), as it defines its key physiological role. It is the most conserved of all CheA domains and no substantial size variation or any duplications of this domain were found in our dataset (Fig. 3B).

**The dimerization domain** (**H-kinase_dim)** is a CheA-specific version of the histidine kinase dimerization domain. We found that it is always occurs in combination with the histidine kinase and CheW domains, as was reported previously for a much smaller dataset (17). The dimerization domain is poorly conserved: in many cases, it was not detected by HMMER (39), an automated sequence-to-profile search tool implemented in MiST database, but upon further investigation we were able to identify it using HHpred, a more sensitive profile-to-profile search tool (40), as a match to a Pfam H-kinase_dim profile or to its structural model (PDB ID 6Y1Y) with probabilities above 90%.

The dimerization domain in CheA of model organisms *E. coli* and *B. subtilis* is small (∼60 a.a.). It is formed by two helices (∼ 26-28 a.a. each) connected by a short loop. A recent study revealed an extended dimerization domain in Spirochetes (41). In CheA of *Treponema denticola* additional ∼50 residues extend the length of the dimerization helices by approximately 2 times (PDB ID 6Y1Y). It was suggested that the extended H-kinase_dim domain might enable stronger dimerization of CheA molecules (41).

While most of the analyzed CheAs in our dataset had the classical short (∼60-70 a.a.) H-kinase_dim domain, many sequences contained its extended versions (>70 a.a.). We found the large range of length variation of the extended dimerization domain, with the longest being up to 3 times of the classical one (170 a.a.). These longest dimerization domains were found in CheAs from Chloroflexota and Cyanobacteria belonging to Tfp system. Thus, it appears that this region of CheA is subject to various duplications resulting in substantial elongation of dimerization helices.

#### CheW domain

This domain is found exclusively in chemosensory systems (17). CheW is one of the core domains in CheA, as it was detected in 100% of all CheA sequences (Fig. 3B). Similarly to the histidine kinase domain, CheW domain is highly conserved, and it was detected by HMMER in every CheA sequence from our dataset. However, in contrast to the kinase domain, we detected instances of its duplication: approximately 8% of CheA sequences in our dataset contained two CheW domains (Fig. 3B). Notably, almost all CheAs with two CheW domains belong to F5 class, which is widely distributed in bacteria (Fig. 2). Two CheW domains were also found in some CheA proteins from F1 class and in one CheA from F10 class. This distribution suggests that the appearance of two CheW domains is a result of independent duplication events.

In several CheA sequences (for example in the *Desulfobacteraceae* family), three CheW domains were identified by HMMER. But upon closer inspection, we concluded that only two CheW domains are present in these proteins. A 50 a.a. insertion in the middle of the N-terminal CheW domain splits CheW into two parts thus “misleading” HMMER into identifying two CheW domains. On the other hand, in two organisms from Chloroflexota (e.g. *Thermomicrobium roseum*) CheA proteins of the F1 class contain three true CheW domains, with one of them separated from another two CheW domains by a CheY-like domain (Fig. 3A).

CheA proteins with two CheW domains have been experimentally studied in alphaproteobacteria *Rhodospirillum centenum* (42), *A. brasilense* (32) and *Caulobacter crescentus* (14) (all belong to F5 class); however, none of the studies specifically explored the function of the two CheW domains. A more recent study of the CheA with two CheW domains from *A. caulinodans* (also belongs to F5 class) showed that the strain where CheW2 was deleted together with the C-terminal response regulator receiver domain, was just as defective in their chemotactic ability as the Δ*cheA* mutant, suggesting that both CheW domains are involved in controlling chemotaxis (33). The presence of more than one CheW domain should significantly change the classical arrangement of bacterial signaling arrays (43); structural studies illuminating the contribution of additional CheW domains would be productive.

#### The phosphotransfer domain (Hpt)

The histidine phosphotransfer domain responsible for the phosphorylation of the aspartate residues in response regulator proteins CheY and CheB. This domain was found in 98.3% of the analyzed CheA sequences, thus initially raising a question whether it is truly a core domain of CheA. Upon further investigation, we found that CheA sequences where no Hpt domain was identified fall in two categories. First, in some instances the CheA was missing Hpt domain due to sequencing and/or assembly errors, as these genes were located at the end of the contig. Second, in some cases where Hpt was missing from CheA, we found the Hpt domain encoded as a separate gene in the same operon. Experimental studies showing that the Hpt domain liberated from the rest of the kinase is fully functional (44), and the operon location of Hpt with the rest of CheA strongly suggest a bipartite system, an arrangement which is not uncommon in bacterial signal transduction (45).

A unique case of partitioning CheA functions between two proteins is seen in *Cereibacter* (*Rhodobacter*) *sphaeroides*, which has two of its four *cheA* genes, namely *cheA3* and *cheA4* located in the same operon (Fig. 3) (46). CheA4 consists of only three core domains: dimerization, kinase, and CheW, whereas CheA3 was reported to contain only Hpt and CheW domains separated by a 794 a.a. region with no identifiable domains (47). Both proteins localize with the cytoplasmic chemoreceptor array and act together as a single functional CheA to control the flagellar motor (46, 47). We were intrigued by the lack of domains in this very long, functionally important region of CheA3 and performed sensitive profile-profile searches using HHpred that confidently identified the following domains in this region: two more Hpt domains (95.94% and 97.40% probability), a P2 domain (96.98%), H_kinase_dim domain (98.14%) and a HATPase_c (97.94%). Sequence alignment showed that the newly identified histidine kinase domain in CheA3 is missing one full alpha helix and parts of the two helices on either side and it does not contain all the residues necessary for Mg2+ and ATP binding. The Alphafold2 (48) model also predicts the histidine kinase domain with missing helices. Thus, we conclude that it is not functional, which agrees with the published data showing a lack of autophosphorylation by CheA3 (46). The Alphafold2 model also shows three Hpt domains. However, in contrast to the first Hpt domain, which is structurally intact, contains a conserved histidine (in position corresponding to His-48 in *E. coli* CheA) and was experimentally shown to be phosphorylated *in vitro* (49), the second and third Hpt domains have structural deviations and lack histidine in a conserved position. Structural deviations are also seen in the P2 domain of CheA3. Taken together, these observations suggest that CheA3 contains structurally modified and largely nonfunctional domains between its N-terminal Hpt and C-terminal CheW domain. It is likely that initially CheA3 was fully functional, but upon duplication of its core region giving the birth to CheA4, corresponding domains of CheA3 lost their function while maintaining their basic structure. These findings also provide an additional explanation why CheA3 and CheA4 work together as a single unit. Twenty other genomes within the *Rhodobacteraceae* family have CheA3 and CheA4 orthologs in similar gene neighborhoods indicating the timing of emergence of this unique CheA system.

CheA homologs with multiple Hpt domains were originally described in chemosensory systems controlling twitching motility in *Synechocystis* PCC6803 (50) and *Pseudomonas aeruginosa* (51). Several homologous CheA proteins with multiple Hpt domains were subsequently identified in genomic studies (17, 21). In our dataset, 18% of CheA sequences contained two or more Hpt domains. Interestingly, more than 90% of those also contained a response regulator receiver domain at the C-terminus. Approximately 93% of the CheA proteins with multiple Hpt domains belong to the ACF and Tfp classes. More than a half of the CheAs with multiple Hpt domains contained five or more copies, with the maximum number (fourteen) detected in the Tfp CheA of a gammaproteobacterium *Oleiphilus messinensis* (Fig. 3A).

The defining feature of the Hpt domain is a conserved histidine residue, which serves as a phosphorylation site, however, some of the Hpt domains in multi-Hpt CheA proteins lack this site. For example, ChpA (CheA of Tfp class) from *Pseudomonas aeruginosa* was reported to have a total of eight Hpt domains, with the conserved histidine only present in six of them (34). The phosphoryl group flow analysis showed that *in vitro* there was no phosphorylation of Hpt domains lacking the conserved histidine and the function of these domains is yet to be established (34). Our analysis revealed that ChpA has one additional Hpt domain without a conserved histidine (Fig. 5). Examination of the ChpA structure predicted by the AlphaFold (UniProt ID Q9I696), revealed that the newly identified Hpt domain consists of four helices (instead of classical five), similarly to the C-terminal Hpt domain, which does contain a conserved histidine (Fig. 5). Additionally, between Hpt1 and newly identified Hpt2 AlphaFold predicted another domain, which structurally resembles Hpt, but it is not recognized as such by HHpred. The role of Hpt domains lacking a conserved phosphorylation site is yet to be determined.

**Fig. 4.**
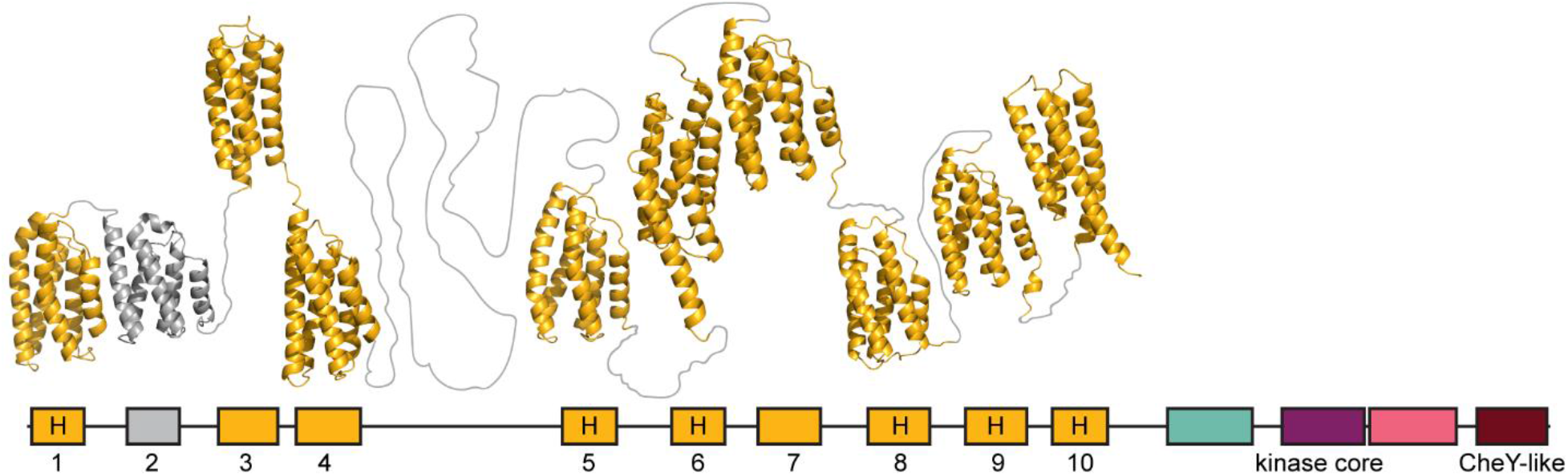
Hpt domains in the ChpA (Tfp CheA) from *P. aeruginosa* PAO1. Domains are colored as in previous figures. Known and predicted Hpt domains are numbered. Domains 2 and 3 were identified in this study. The presence of a conserved histidine residue, corresponding to His-48 in *E. coli* CheA is marked by “H”. Structure is predicted by AlphaFold (48).

**Fig. 5.**
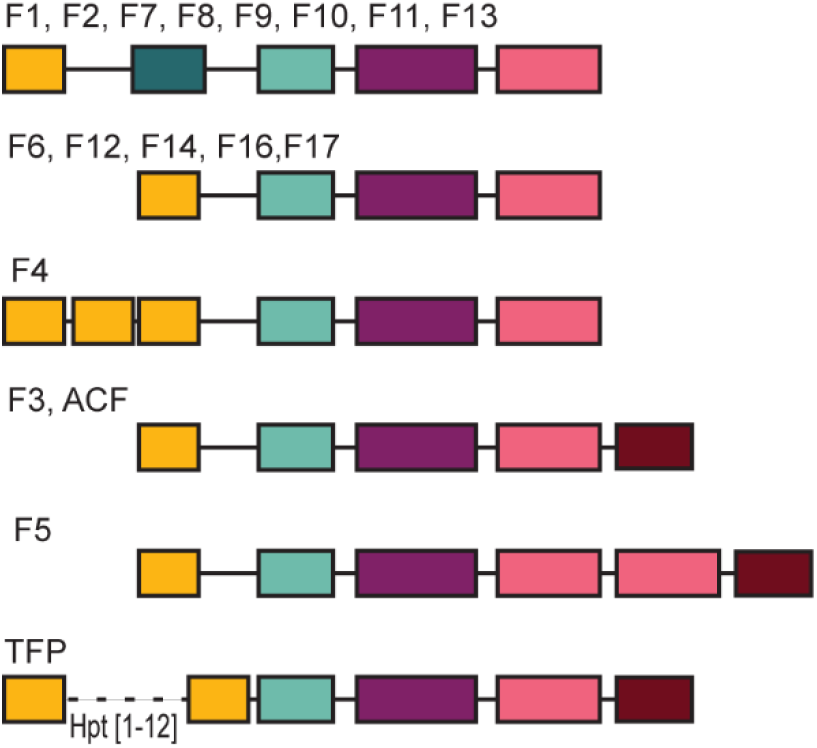
A predominant domain composition of CheA proteins from different chemosensory classes. Chemosensory classes are defined in (17). Domains are colored as in previous figures.

### Auxiliary domains of CheA

While the four core domains are the minimum requirement for CheA function, many CheA proteins contain additional domains. For example, the classical five-domain CheA protein in *E. coli* has a P2 domain (Fig. 1), which promotes CheY binding.

#### P2/CheY-binding

The Pfam database has two domain models, P2 and CheY-binding, that belong to the same clan (Table S1). The existence of two models for this domain is likely due to substantial sequence variability: this domain is much less conserved than any of the CheA core domains (21). The P2 model was developed using CheA from *T. maritima*, the model organism for structural biology, and related organisms (e.g. *B. subtilis*), whereas the CheY-binding model was developed using CheA from *E. coli*, where its role was studied experimentally, and related organisms. However, structurally, these versions of the P2 domain match closely (26), thus we refer to this domain as P2/CheY-binding. Initially, this domain was identified in less than 20% of all CheA sequences in our dataset (detected by HMMER as P2 in 1,960 sequences and as CheY-binding in 873 sequences). We then used sensitive profile-profile searches with HHpred and identified a total of 6,519 sequences with P2/CheY-binding domains, which means that more than 50% of CheA sequences in our dataset lack this domain (Fig. 3B). The lack of P2/CheY-binding domain is further supported by AlphaFold models that show an extended unstructured region between the Hpt and the dimerization domain in sequences where no P2/CheY-binding domain was identified by HHpred. On one hand, this was a surprising finding, because best studied CheA proteins from model organisms *E. coli, S. enterica, B. subtilis,* and *T. maritima* all contained this domain. On the other hand, it was known that CheA proteins in *C. jejuni* and *H. pylori* do not have this domain, and even in *E. coli* it is not essential for the key CheA function – phosphotransfer to CheY (52). Deletion of P2 results in much slower phosphotransfer rates and therefore impaired chemotaxis ability (20, 52); however, overexpression of Hpt domain could correct for the lack of a P2 domain (20). Thus, the fact that in nature the P2/CheY-binding domain is dispensable may not be surprising after all.

The vast majority of CheA proteins contain only one copy of the P2/CheY-binding domain, although we detected some sequences with duplications of this domain (Fig. 3). Notably, duplication of this domain occurred in the common ancestor of *Borreliales* (Spirochaetota phylum), as all members of this order have CheA proteins with a duplicated P2 domain. Phyletic distribution of other CheA proteins with a duplicated P2/CheY-binding domain suggests several independent duplication events.

### Response regulator receiver domain (CheY-like)

Together with the histidine kinase domain, the response regulator receiver domain comprises the essential core of bacterial two-component signal transduction systems (53). Current domain databases define it as a superfamily (Pfam accession CL0304 termed CheY-like) containing several families, one of which has a more general name of a response regulator receiver domain (Pfam accession PF00072 termed Response_reg). In model chemosensory systems from *E. coli* and *B. subtilis*, this domain is present in two response regulators – CheY and CheB. However, in other homologous systems, for example in *C. jejuni* and *H. pylori*, it was also found as a component of CheA. The function of the CheY-like domain in CheA homologs has been studied experimentally in several organisms, and its common role appears to be that of the phosphate sink (54), as shown in *H. pylori* (55). In *P. aeruginosa* ChpA, the CheY-like domain was shown to potentially function as a phosphate sink and/or a source of phosphoryl groups for two of the Hpt domains that do not have a conserved histidine residue (34). In *A. caulinodans* CheA, the CheY-like domain does not seem to function as a phosphate sink but it is necessary for the dephosphorylation of the Hpt domain (33). In other systems, the role of this domain has not been studied in detail, but it is known to be important for the proper function of the CheA kinase (32, 33, 56, 57). For example, the disruption of the CheY-like domain of *R. centenum* CheA (CheA1) caused the lack of chemotaxis and phototaxis (33, 57).

We have identified the CheY-like domain in approximately 34% of analyzed CheA sequences (Fig. 3). The majority of those belong to four CheA classes: F3, F5, ACF, and Tfp. Additionally, several sequences belonged to F1 and F7, whereas the rest of CheAs from all other flagellar classes did not have this domain. In most cases, the CheY-like domain was at the CheA C-terminus, however, we found several cases where this domain was located at the N-terminus (Fig. 3A). While the majority of the CheAs sequences contained only one CheY-like domain (Fig. 3B), some had two, and two sequences, including the Tfp class CheA from a cyanobacterium *Spirulina major,* contained three CheY-like domains (Fig. 3B).

To test a hypothesis that the main function of the CheY-like domain is the role of a phosphate sink, as shown for *H. pylori* CheA (55), we analyzed the presence of dedicated phosphatases CheZ (10), CheC and CheX (58) in genomes that had a single CheA protein containing the CheY-like domain. Most of bacterial chemosensory systems employ either CheZ or CheC/CheX type phosphatases (17). Thus, we argue that the absence of phosphatase genes in genomes containing a single CheA with the CheY-like domain would support the phosphate sink role for this domain. Satisfactorily, we found that 93% of such genomes (811 out of 871) lacked chemosensory phosphatases.

#### CheC/CheX

CheC and CheX are protein phosphatases that dephosphorylate CheY (58). Both phosphatases are part of the Pfam CheC-like clan (CL0355) and have similar structural topologies (58). In our dataset, we found CheC/CheX fusion with F1 CheAs in *Leptospirae* class of spirochetes. Given that these domains dephosphorylate CheY, it is possible that such CheA-CheC/CheX fusions have a dual kinase/phosphatase function and are able to phosphorylate and dephosphorylate CheY.

### Domain co-occurrence and class-specific domain composition

In addition to invariable co-occurrence of the four core domains, we noticed the following trends. First, more than 90% of CheA protein sequences containing multiple Hpt domains also had a CheY-like domain. This trend is predominant among Tfp class (94%) but it is also found in ACF class. Second, the majority of CheA homologs that contain the CheY-like domain do not have the P2/CheY-binding domain. Finally, domain architectures are generally class-specific (Fig. 6). For example, all CheA sequences from F4 class contain three Hpt domains and 99% of CheA sequences from F5 class have a duplicated CheW domain.

## Conclusions

CheA homologs have four core domains: (i) the phosphotransfer domain, which can be present in multiple copies and, occasionally, reside as a separate gene in the same gene neighborhood, (ii) the dimerization domain of a variable length, (iii) the histidine kinase domain, which is always present in a single copy, and (iv) the CheW domain, which is duplicated in some homologs. The P2/CheY-binding domain, which enhances phosphotransfer, and the CheY-like domain, which likely serves as a phosphate sink, are the most common auxiliary domains found in CheA homologs. CheA homologs from different classes have a predominant domain composition.

In spite of their high specificity, current models (profile HMMs) for the Hpt, P2/ CheY-binding, and dimerization domains models have low sensitivity and perform poorly in automated sequence similarity searches, often resulting in missing domains. However, models for the histidine kinase (HATPase_c) and CheW domains are both highly sensitive and specific. Thus, because the presence of these two core domains uniquely distinguishes CheA homologs from other proteins, CheA sequences can be easily identified by automated searches and then further explored for the presence of other domains using more sensitive approaches, such as. HHpred (40) and AlphaFold (48).

## Materials and Methods

The following databases were used in this study: GTDB release R95 (36), MiST v3.0 (38), AnnoTree v1.3 (59).

The domain architectures for all collected CheA sequences were identified using two approaches. First, domains were predicted using HMMER (39). Then, in cases where no domain was identified by HMMER in sequence regions longer than 100 amino acid residues (the average domain size), we ran a more sensitive domain identification search tool HHpred (40), using the region of interest as a query. All sequence similarity searches were performed with default parameters. Archaeal CheA sequences were gathered from AnnoTree hits using CheW domain as a query and subtracting CheW protein sequences. Their domain composition and class were predicted using TREND (60, 61).

Multiple sequence alignments were built using MAFFT (62) with automatically selected parameters; structural models were built by AlphaFold2 (48).

## Acknowledgements

This work was supported by the National Institutes of Health R35GM131760 (to I.B.Z.).

